# Identification of Differentially Expressed Genes and Proteins Related to Diapause in *Lymantria dispar*: Insights for the Mechanism of Diapause from Transcriptome and Proteome Analyses

**DOI:** 10.1101/2024.12.07.627326

**Authors:** Qing Xie, Xiaofan Ma, Yafei Li, Wenzhuai Ji, Fengrui Dou, Xiue Zhu, Juan Shi, Yixia Cao

## Abstract

Spongy moth (*Lymantria dispar* Linnaeus) is a globally recognized quarantine leaf-eating pest. Spongy moths typically enter diapause after completing embryonic development and overwinter in the egg stage. They spend three-quarters of their life cycle (approximately nine months) in the egg stage, which requires a period of low-temperature stimulation to break diapause and continue growth and development. In this study, we explored the molecular mechanism underlying the diapause process in spongy moth. We performed bioinformatics analysis on four Asian populations of spongy moth and one Asian–European hybrid population through a transcriptome analysis combined with proteomics. The results revealed that 1,842 genes were differentially expressed upon diapause initiation, while 264 genes were identified upon diapause termination. Eight diapause-related genes were screened out from the three-level pathways that were significantly enriched by differentially expressed genes at the time of diapause and diapause termination, and the phylogenetic tree and protein three-dimensional structure model were constructed. This study elucidates the diapause mechanism of spongy moth at the gene and protein levels, providing theoretical insights into the early and precise prevention and control of spongy moth. This study can facilitate the development of an efficient, environmentally friendly control system for managing spongy moth populations in the field.

## 1. Introduction

The spongy moth (*Lymantria dispar* Linnaeus, 1758), also known as the gypsy moth, originates from the Eurasian continent. Currently, the spongy moth has spread to over 50 countries and regions across Europe, the Americas, Africa, and Asia. Being a well-known forest pest in the Holarctic region [1], the spongy moths are characterized by their polyphagous feeding habits, strong dispersal ability, and the tendency to cause severe damage. This pest has caused numerous outbreaks [2], and its larvae can feed on 300–500 species of broadleaf trees and several species of conifers [3, 4], causing substantial ecological and socioeconomic losses. Thus, this forest pest requires focused control measures.

Spongy moths can be categorized into three subspecies on the basis of the flight capacity of female moths and their geographical spread: the European spongy moth (ESM), the Asian subspecies, and the Japanese subspecies. Collectively, they are referred to as the flighted spongy moth complex (FSMC) [5]. A key difference among these subspecies is that female moths of the ESM cannot fly upward [6, 7], unlike those in the FSMC that can sustain upward flight [8–10]. Moreover, the FSMC requires a shorter period for diapause termination at low temperatures than the ESM [11].

During their growth, development, and reproduction, insects encounter adverse environmental conditions, such as low temperatures. To cope with adversities, they may employ self-regulation mechanisms to halt their growth and development at a certain stage, thus resisting the adverse environmental effects and ensuring the continuity of their population. This adaptive strategy, known as diapause [12], can be classified into two types: obligate diapause, which occurs at fixed generations and life stages, and facultative diapause, which is regulated by external environmental factors, such as photoperiod, temperature, and food availability. In particular, the spongy moths undergo obligate diapause, entering this phase after completing embryonic development and overwintering as eggs. Thus, three-quarters of the spongy moths’ annual life cycle (approximately nine months) are spent in the egg stage.

Factors affecting insect diapause include external environmental conditions and internal hormonal changes. Temperature and photoperiod are two major environmental factors affecting insect diapause. In 1991, Gray et al. used the Li6200 photosynthesis system to measure the respiratory rate of ESM eggs under various temperatures. They observed that the eggs maintained a low respiratory rate at 4°C, exhibiting an initial increase and then a decrease from 0 to 40 days after oviposition under all other temperature conditions, demonstrating substantial temperature effects on the respiratory rate of the eggs [13]. Another study confirmed that the eggs of spongy moths do not hatch unless exposed to cold treatment for adequate durations following embryonic development [14], indicating the necessity for cold stimulation to terminate diapause and enable growth and development. In 2014, Li analyzed the hatching rates of FSMC larvae under various photoperiods. Their results revealed that photoperiod had no considerable effect on diapause termination in spongy moths [14].

In subsequent studies, Gray et al. determined that the eggs of ESM gradually enter a diapause induction phase 15 days after the completion of embryonic development, reaching a complete diapause stage at day 40 after oviposition. These findings indicated that ESM exhibited the shortest duration to break diapause when transferred to a 25°C environment after 75 days of exposure to 5°C [15]. Moreover, the diapause of spongy moth eggs was categorized into the pre-diapause, diapause, and post-diapause stages, collectively termed the diapause program by subsequent researchers [16]. In 2012, Wei conducted experiments by using a completely randomized and second-order orthogonal regression design, confirming that the diapause of FSMC could be broken after 60 days of cold stimulation at 5°C [17]. This finding indicates that compared with ESM, the FSMC has a weaker diapause intensity, thus requiring a shorter time for diapause termination. This observation has been confirmed in subsequent studies [11, 18, 19].

In addition to environmental factors that affect diapause in spongy moths, internal hormones play a regulatory role during diapause. Lee et al. determined that ecdysteroids, such as ecdysone, are involved in the diapause of spongy moths, with a decrease in ecdysteroid titers being closely associated with diapause termination [20]. However, Atay-Kadiri et al. reported that the application of the exogenous hormone ecdysone had no effect on diapause in spongy moths [21]. A study on the FSMC revealed that trehalose, glucose, and polyol small molecules, such as glycerol, mannitol, sorbitol, and inositol, play essential roles in diapause and response to cold stimulation in the spongy moths [22].

Current transcriptomic and proteomic studies on diapause processes have been conducted in organisms such as *Bombyx mori* [23–25], *Helicoverpa armigera* [26–28], *Leptinotarsa decemlineata* [29], and *Chilo suppressalis* [30]. However, studies on diapause in the spongy moths have mainly investigated the effects of external environmental factors and metabolic substances, without a systematic exploration at the molecular level. Thus, this study focused on four FSMC populations and one Eurasian hybrid population. We used transcriptomic methods, including mRNA sequencing, and proteomic methods, such as tandem mass tag (TMT) labeling, to perform bioinformatics analysis, screen for diapause-associated genes, and explore the differential expression patterns of genes and proteins throughout the diapause process in the spongy moths.

## 2. Materials and Methods

### 2.1 Insect Strains

The egg masses from the Inner Mongolia (NMG), Shanxi (SX), Liaoning (LN), and Yunnan (YN) populations of *L. dispar* asiatica (FSMC) were collected in the wild from host trees, raised in the Plant Quarantine Laboratory of Beijing Forestry University. In the laboratory, they were reared on an artificial diet until pupation. Upon eclosion, individuals from the same geographical population were paired for mating. In addition, a Eurasian hybrid strain, ONMG, was created by mating male moths from the New Jersey (NJ) population of the United States with female moths from the NMG population of China in the laboratory. The egg masses resulting from these matings were used in the subsequent experiment. Table 1 provides specific information.

**Table 1.** Location information of the sampling sites for the spongy moth, *Lymantria dispar*.

| Population Code | Sampling Location | Latitude | Longitude |
| --- | --- | --- | --- |
| NMG | Chifeng, Inner Mongolia, China | 41°38'N | 118°53'E |
| SX | Shuozhou, Shanxi, China | 39°23'N | 112°66'E |
| LN | Dalian, Liaoning, China | 38°56'N | 121°34'E |
| YN | Fugong, Yunnan, China | 27°09'N | 98°52'E |
| NJ | New Jersey, USA (New Jersey Standard Strain) | 39°41'N | 75°45'W |

During the pre-diapause, diapause, and post-diapause periods, egg masses laid at the same time were selected for the experiment. The division of diapause stages was based on the findings reported by Wang [31]. The egg masses from each population during these three periods were respectively named using the abbreviation of the population followed by 1, 2, or 3 (e.g., pre-diapause egg masses of SX were labeled as “SX1”). Each experimental group comprised 500 eggs from each test population, and three replicates were used in the transcriptomic and proteomic analyses. The sampled egg masses were stored at −80°C in a freezer for subsequent experiments.

### 2.2 Transcriptome Sequencing and Differential Expression Analysis of Spongy Moths at Different Diapause Stages

#### 2.2.1 RNA Extraction, Quality Control, Library Construction, and Sequencing

The sampled egg masses were thoroughly homogenized in lysis buffer, and the supernatant was collected. RNA was extracted using the TRIzol reagent kit (Invitrogen, Carlsbad, CA, USA), and the purity and concentration of RNA were determined using a Nanodrop 2000 UV spectrophotometer. The quality and integrity of RNA were evaluated using agarose gel electrophoresis and the Agilent 2100 Bioanalyzer (Agilent Technologies, Palo Alto, CA, USA). mRNA was enriched using oligo(dT) magnetic beads, followed by fragmentation with fragmentation buffer to obtain fragments approximately 300 bp in size. Single-stranded cDNA was synthesized from mRNA templates by using random hexamers in the presence of reverse transcriptase, followed by second-strand synthesis to form stable double-stranded structures. The sticky ends of double-stranded cDNA were repaired to obtain blunt ends by using the end repair mix, and an “A” base was added to the 3’ end for adapter ligation. After adapter ligation, the products were purified and selected based on their size. The size-selected products were subjected to PCR amplification and purification, and subsequently sequenced on the Illumina NovaSeq 6000 sequencing platform (Shanghai Majorbio Bio-pharm Technology Co., Ltd., Shanghai, China).

#### 2.2.2 Transcriptome De Novo Assembly and Functional Annotation

To ensure the quality of subsequent bioinformatics analyses, we systematically assessed the base distribution and quality fluctuations of reads from all sequencing runs as indicators of sequencing and library construction quality. This included statistical analyses of base composition distribution, base error rate distribution, and base quality distribution for each sequencing cycle. Subsequently, raw sequencing data were filtered to obtain clean data, facilitating downstream analysis for de novo transcriptome studies in the absence of a reference genome. High-quality RNA-seq sequencing data acquisition was followed by de novo assembly to generate singleton sequences and contigs. This process yielded all transcripts from the current transcriptome sequencing experiment.

The obtained transcripts were subjected to the following analyses:

1. Functional annotation: Transcripts were compared against six major databases (i.e., Nonredundant [NR], Swiss-Prot, Pfam, Clusters of Orthologous Groups [COG], Gene Ontology [GO], and Kyoto Encyclopedia of Genes and Genomes [KEGG]) to obtain annotation information for the transcripts and genes;
2. Expression level analysis: RNA-Seq by Expectation-Maximization (RSEM) was performed to quantitatively analyze the expression levels of genes and transcripts;
3. Differential expression analysis: DESeq2 was used to analyze and identify differential expression of genes based on read counts between samples;
4. Enrichment analysis of differential genes in GO and KEGG pathways: GO enrichment analysis was performed using Goatools, and Fisher’s exact test was used for precise examination. GO categories with corrected p values (pfdr) ≤ 0.05 were considered significantly enriched. KEGG pathway enrichment analysis was conducted using KEGG Orthology Based Annotation System (KOBAS), and Fisher’s exact test was performed for precise examination. The Benjamini–Hochberg (BH) method was used for multiple testing correction to control the false discovery rate. KEGG pathways with corrected p values ≤ 0.05 were considered significantly enriched among differentially expressed genes, from which genes associated with diapause were selected.

#### 2.2.3 Homology and Phylogenetic Analysis of Diapause-Associated Genes in Spongy Moths

Homology of diapause-associated genes in spongy moths was examined by aligning sequences against the NCBI database (https://blast.ncbi.nlm.nih.gov/Blast), and amino acid sequences with a similarity of >60% were downloaded for phylogenetic analysis. The neighbor-joining (NJ) method was used in MEGA 7.0 to construct phylogenetic trees, with confidence assessed through 1000 bootstrap replicates.

#### 2.2.4 Construction of Three-Dimensional Models of Diapause-Associated Genes

The amino acid sequences of diapause-associated genes in spongy moths were aligned on the SwissModel website (https://swissmodel.expasy.org/) to generate protein structure predictions. Models with the highest global model quality estimation values (all exceeding 0.90) were selected as reference templates. Subsequently, protein secondary structures were predicted using the Jpred4 website (https://www.compbio.dundee.ac.uk/jpred4/index_up.html) to enhance the accuracy of protein structure predictions.

At the MolProbity website (http://molprobity.biochem.duke.edu/) and SwissModel website, all three-dimensional models of diapause-associated proteins were evaluated. The Ramachandran plot was used to illustrate whether the dihedral angles of amino acid residues in the main chain of the protein were within reasonable ranges. The MolProbity index was used to evaluate the quality of all atoms within a conformation, including side-chain atoms. MolProbity scoring consider atomic contacts, clashes, bond lengths, angles, and torsion angles.

### 2.3 Proteomic Analysis of Spongy Moths Using Tandem Mass Tag Labeling

#### 2.3.1 Protein Extraction

Samples stored at −80°C were ground into powder in liquid nitrogen. Subsequently, each 100 mg of the sample was dissolved in 1 mL of lysis buffer. After sonication for 5 min to remove nucleic acids, the mixture was centrifuged at 3000 rpm for 20 min at 4°C, and the precipitate was collected. The precipitate was then re-suspended in 800 μL of cold acetone to remove impurities. After centrifugation at 3000 rpm for 20 min at 4°C, the precipitate was collected and air-dried. Finally, 600 μL of 8 M urea was added to dissolve the proteins, and the total protein sample was obtained. The protein sample concentration was determined using the Bradford method [32].

#### 2.3.2 SDS-PAGE Electrophoresis

We extracted 30 μg of the protein sample and diluted it to a final volume of 15 μL. Then, 5 μL of buffer was added, and the mixture was incubated at 95°C water bath for 5–10 min, followed by cooling to room temperature in an ice-water bath and brief centrifugation. The electrophoresis chamber, equipped with a glass plate assembly, was filled with electrode buffer. Approximately 1/4 to 1/3 of the chamber was filled with the same buffer. Then, 20 μL of protein solution was loaded into the sample wells below the buffer. Sequential electrophoresis runs were performed, including a 20-min run at 80 V for the stacking gel, followed by a 60-min run at 150 V for the resolving gel.

#### 2.3.3 Reduction, Alkylation, and Enzymatic Digestion

Triethylammonium bicarbonate buffer (TEAB) was added to the protein sample to achieve a final concentration of 100 mM per 100 μg of protein. Then, tri(2- carboxyethyl)phosphine was added to reach a final concentration of 10 mM, and the mixture was incubated for 60 min. Subsequently, iodoacetamide (IAM) was added to a final concentration of 40 mM, and the reaction was conducted for 40 min in dark conditions. Acetone was then added, and the mixture was precipitated at −20°C for 4 h. After centrifugation at 14,000 rpm for 20 min, the precipitate was collected, and 100 µL of 100 mM TEAB was added to the precipitate. The mixture was then allowed to dissolve completely. Trypsin was added to the solution at an enzyme-to-protein ratio of 1:50, and the mixture was digested at 37°C for at least 12 h.

#### 2.3.4 Tandem Mass Tag Labeling

Tandem mass tag (TMT) label reagent (catalog number 90111, ThermoFisher Scientific, Waltham, MA, USA) was mixed with acetonitrile, and the mixture was vortexed and then centrifuged. Then, 100 μg of the aforementioned TMT reagent was added to each sample tube and incubated at room temperature for 2 h. Hydroxylamine was added, and the mixture was incubated for 30 min at room temperature. Finally, the labeled products of each group were obtained and vacuum freeze-dried for later use.

#### 2.3.5 Ultra-performance Liquid Chromatography Gradient Fractionation

We used the high-pH reversed-phase ultra-performance liquid chromatography (UPLC) fractionation system equipped with the ACQUITY UPLC BEH C18 column to fractionate peptide segments. Phase A consists of 2% acetonitrile (pH 10), whereas phase B consists of 80% acetonitrile (pH 10), with a UV detection wavelength set at 214 nm. The UPLC gradient was programmed as follows: flow rate of 200 μL/min, the gradient started at 0%–5% B for elution from 0 to 17 min, followed by 5%–10% B from 17 to 18 min, 10%–30% B from 18 to 35.5 min, 30%–36% B from 35.5 to 38 min, 36%–42% B from 38 to 39 min, 42%–100% B from 39 to 40 min, and maintained at 100% B until 48 min. The collected 28 fractions were pooled into 14 fractions and then dissolved in a mixture of 2% acetonitrile and 0.1% formic acid buffer solution after negative pressure centrifugation for secondary analysis.

#### 2.3.6 Liquid Chromatography–Mass Spectrometry Analysis

Using the EASY-nLC liquid chromatography system, peptide segments were separated using mobile phase A (0.1% formic acid) and mobile phase B (100% acetonitrile with 0.1% formic acid).

The liquid chromatography elution gradient was as follows: at a flow rate of 300 nL/min, the gradient started at 5%–38% B for elution from 0 to 30 min, followed by 38%–90% B from 38 to 90 min, and maintained at 90% B until 44 min.

The separated peptide segments were analyzed using the Orbitrap Exploris 480 mass spectrometry system (Thermo, USA) in the data-dependent acquisition mode, with an MS scan range set at 350–1500 m/z. Data acquisition and analysis were performed using Thermo Xcalibur 4.0.

#### 2.3.7 Database Search

The raw data were analyzed for protein peptide matching and database search identification by using Proteome Discoverer TMSoftware 2.4 software. The database used was all_predicted.pep_unique.fasta. The search parameters were set as follows: trypsin digestion, with up to two missed cleavage sites allowed, a primary mass tolerance of 20 ppm, and a secondary mass tolerance of 0.02 Da.

## 3. Results

### 3.1 Transcriptome Sequencing and Identification of Proteins in Spongy Moths

To construct cDNA sequencing libraries, we used total RNA extracted from the egg samples collected at the pre-diapause, diapause, and post-diapause stages of spongy moths from five geographic populations. These libraries were sequenced on the Illumina HiSeq platform, resulting in 5.79 Gb of clean data with a Q30 base ratio of 92.96%. The de novo assembly of all clean data from the samples was performed using the software Trinity. The assembled unigenes and transcripts were then merged with all the clean data, leading to mapping rates of 77.99% and 85.64%, respectively. A total of 72,812 unigenes and 100,564 transcripts were obtained, with an average N50 length of 1799 bp. Unigenes accounted for 72.40% of the total transcripts. The length distribution of unigenes (Fig 1A) ranged from 201 to 31,697 bp, with an average sequence length of 933 bp. The majority of the unigenes had lengths ranging from 200 to 500 bp, accounting for 55% of the total count.

**Fig 1.**
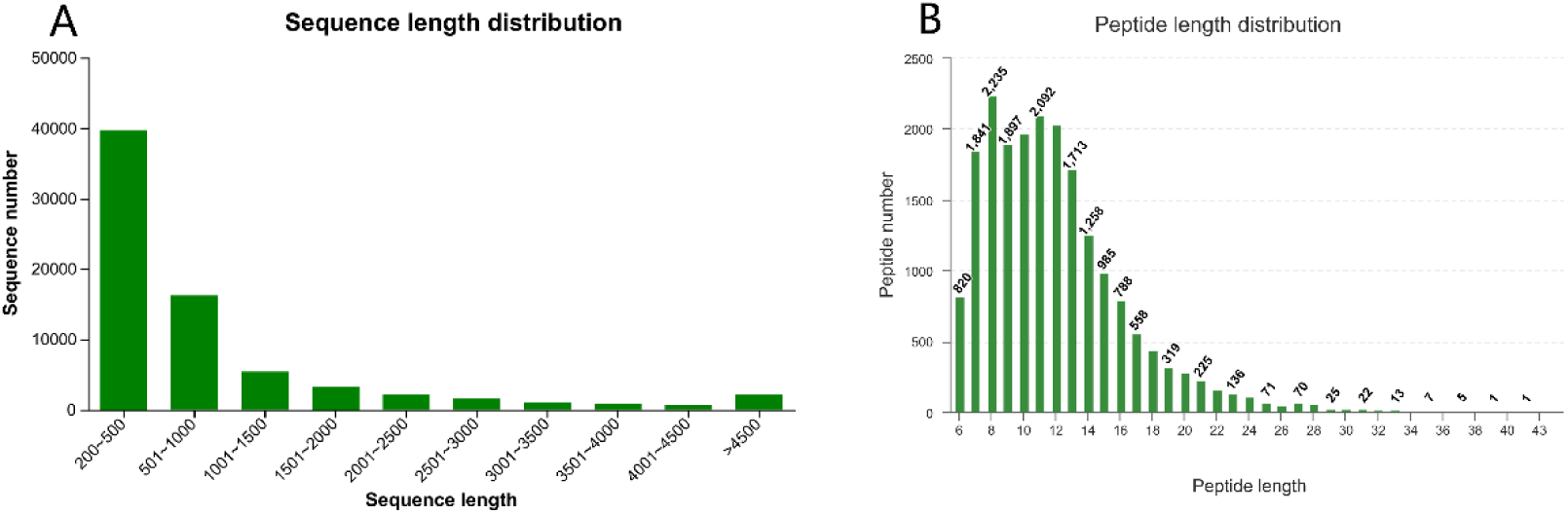
Distribution of sequence length (A) and peptide length (B).

For protein identification, 191,240 total spectra were processed, resulting in 26,397 identified spectra. From these, 20,235 peptides were identified, corresponding to 7,644 proteins and 4,499 protein groups. The distribution of peptide lengths is presented in Fig 1B.

### 3.2 Functional Annotation

We performed transcriptomic analysis without a reference genome. The assembled unigenes were annotated using six major databases: NR, Swiss-Prot, Pfam, eggNOG, GO, and KEGG (Table 2). Among these unigenes, 5,708 were successfully annotated across all databases, whereas 26,435 were annotated in at least one database. See data file S1-S7 in the Supplementary Material for detailed information.

**Table 2.** Transcriptome and protein annotation statistics of *Lymantria dispar*.

| Database | Number of Unigenes | Proportion (%) | Protein Number | Proportion (%) |
| --- | --- | --- | --- | --- |
| NR | 26003 | 35.99 | - | - |
| Swiss-prot | 12448 | 17.23 | - | - |
| Pfam | 13657 | 18.90 | 3994 | 88.78 |
| eggNOG | 20273 | 28.06 | 4372 | 97.18 |
| GO | 14554 | 20.14 | 2658 | 59.08 |
| KEGG | 10288 | 14.24 | 3131 | 69.59 |
| SubCell-Location | - | - | 4499 | 100 |
| Total_anno | 26435 | 36.59 | 4499 | 100 |
| Total | 72255 | 100 | 4499 | 100 |

All proteins were analyzed using mass spectrometry, and their sequences were compared with four major databases (Pfam, eggNOG, GO, and KEGG) and relevant subcellular localization databases (Table 1). All 4,499 proteins were annotated in at least one database, and 1,426 of them were successfully annotated in all databases. See data file S8-S12 in the Supplementary Material for detailed information.

On the basis of the homologous alignment of spongy moth unigenes in the NR database, we constructed a distribution chart of NR annotations for species (Fig 2). The results indicated that the most frequently aligned species was *Arctia plantaginis* (23.57%), followed by *H. armigera* (4.30%), *Trichoplusia ni* (3.97%), *Eumeta japonica* (3.48%), and *Spodoptera frugiperda* (3.07%).

**Fig 2.**
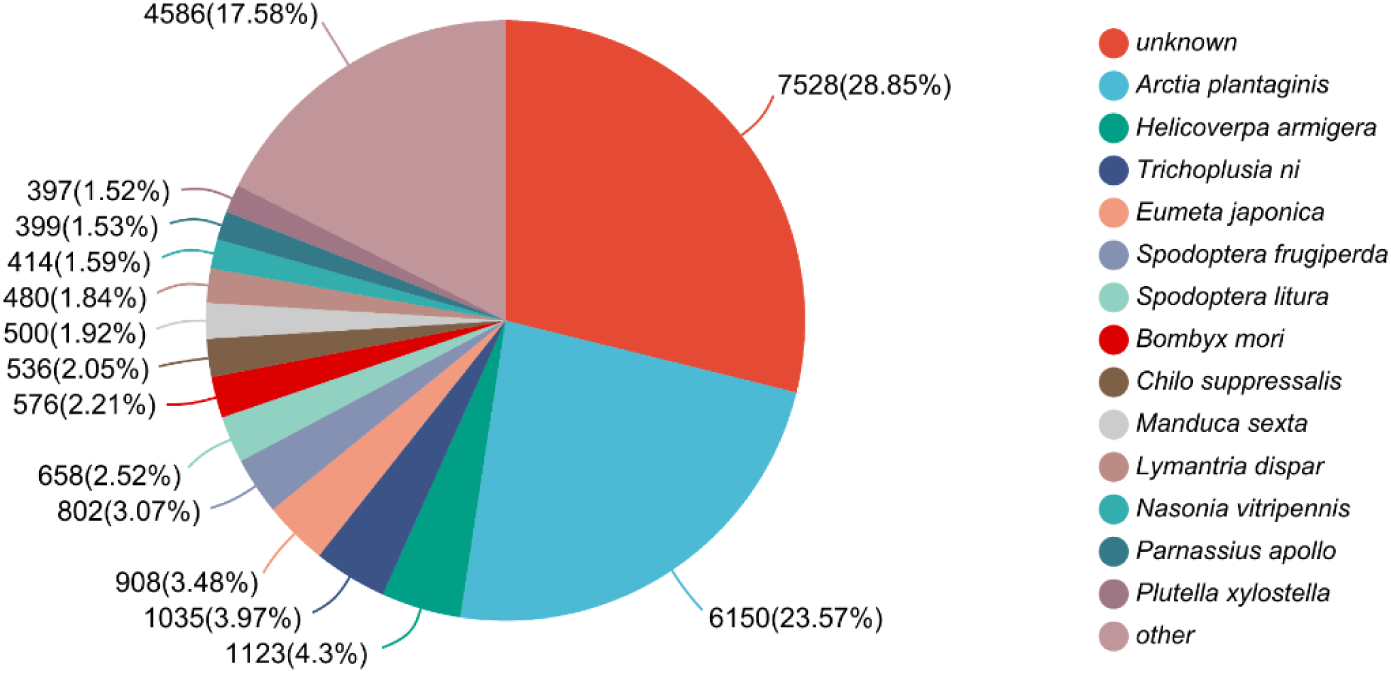
Species distribution of *Lymantria dispar* in the NR database.

A total of 14,554 unigenes and 2,658 proteins from the spongy moths were successfully annotated in the GO database and categorized into three main classes: cellular component (CC), biological process (BP), and molecular function (MF; Fig 3). In the CC category, the cell part had the highest number of unigenes, whereas the cellular anatomical entity had the highest number of proteins. In the BP category, the cellular process entity had the highest count for both unigenes and proteins. In the MF category, catalytic activity had the highest number of unigenes, whereas binding had the highest number of proteins.

**Fig 3.**
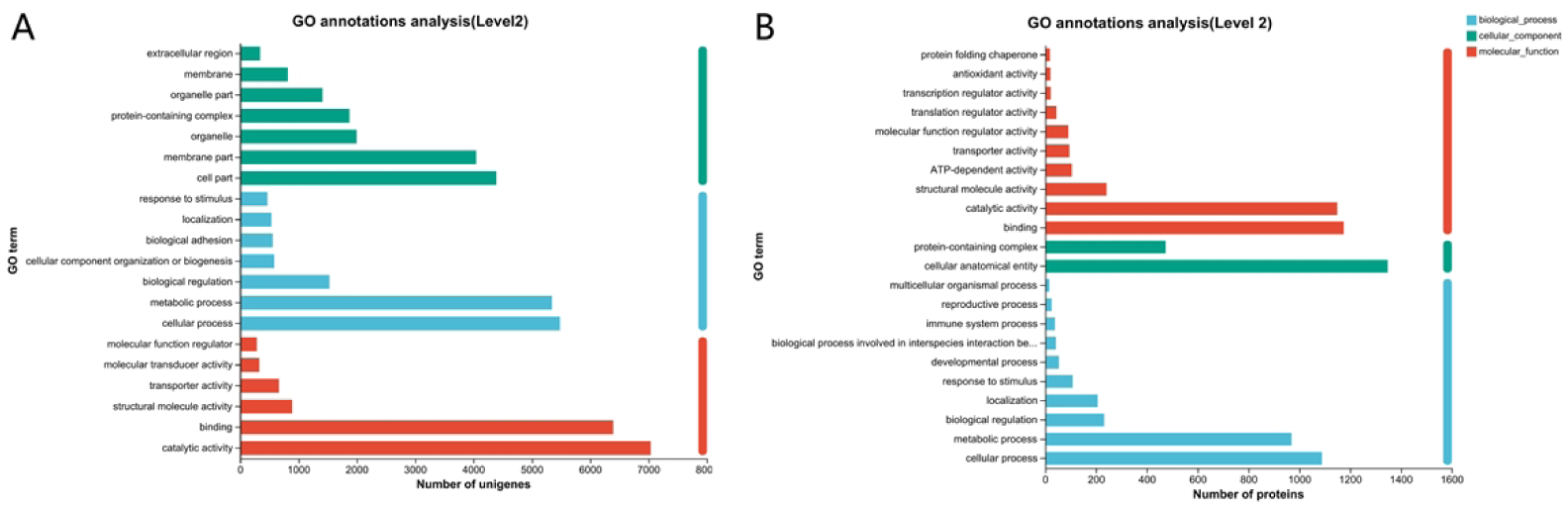
Transcriptome (A) and proteome (B) GO function distribution statistics of *Lymantria dispar*.

In the KEGG database, 10,288 unigenes and 3,131 proteins of the spongy moths were successfully annotated into six metabolic pathways: metabolism, genetic information processing (GIP), environmental information processing (EIP), cellular processes, organismal systems (OS), and human diseases (Fig 4). Among these pathways, except for human diseases, signal transduction had the highest number of unigenes and proteins, followed by transport and catabolism.

**Fig 4.**
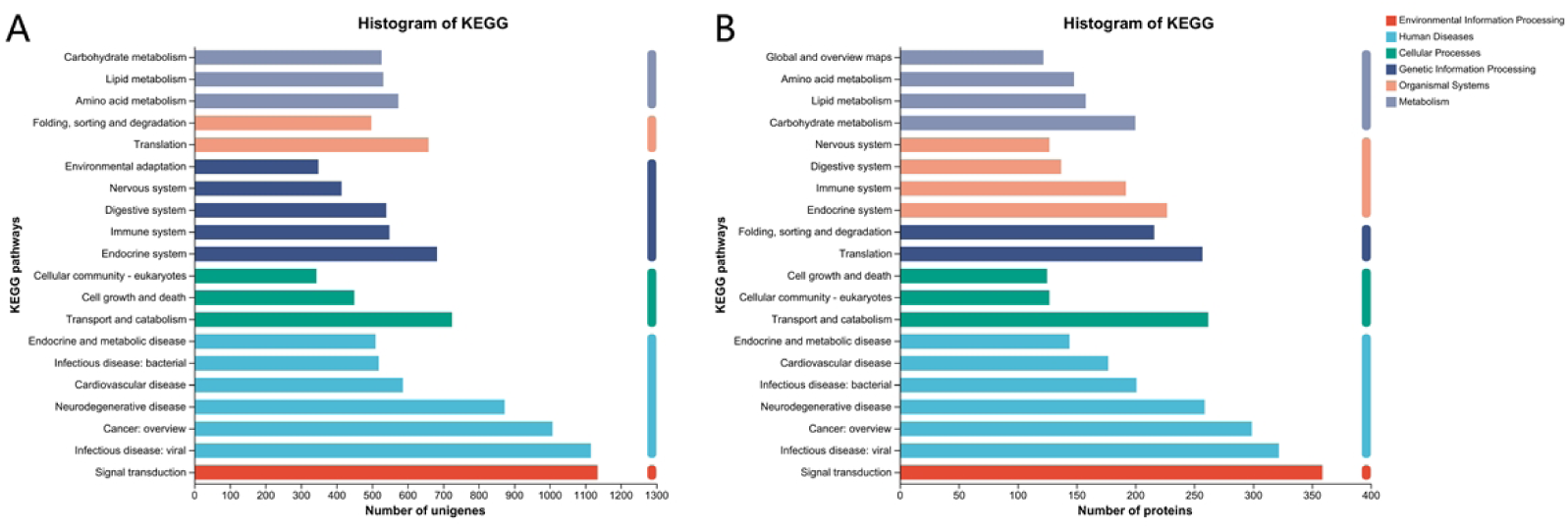
Transcriptome (A) and proteome (B) KEGG pathway distribution statistics of *Lymantria dispar*.

### 3.3 Differential Gene and Protein Expression Analyses

On the basis of differences in expression levels across various diapause stages, sets of proteins and genes were identified for intergroup differential expression analysis. The period from pre-diapause to mid-diapause was termed as the diapause initiation stage, whereas the period from mid-diapause to post-diapause was termed as the diapause termination stage.

During the diapause initiation stage, 1,842 genes were differentially expressed across the five populations, with 470 showing an upregulation and 465 showing a downregulation, while the remaining genes exhibited differential regulation across populations. In addition, 1,325 proteins, with 822 being upregulated and 264 being downregulated, were differentially expressed across the five populations. During the diapause termination stage, 264 genes were differentially expressed across the five populations, which included 87 upregulated and 58 downregulated genes. Furthermore, 442 proteins were differentially expressed among the five populations, including 123 upregulated and 83 downregulated proteins. Fig 5 presents the numbers of differentially expressed genes and proteins at different diapause stages for each population. See data file S13-S16 in the Supplementary Material for detailed information.

**Fig 5.**
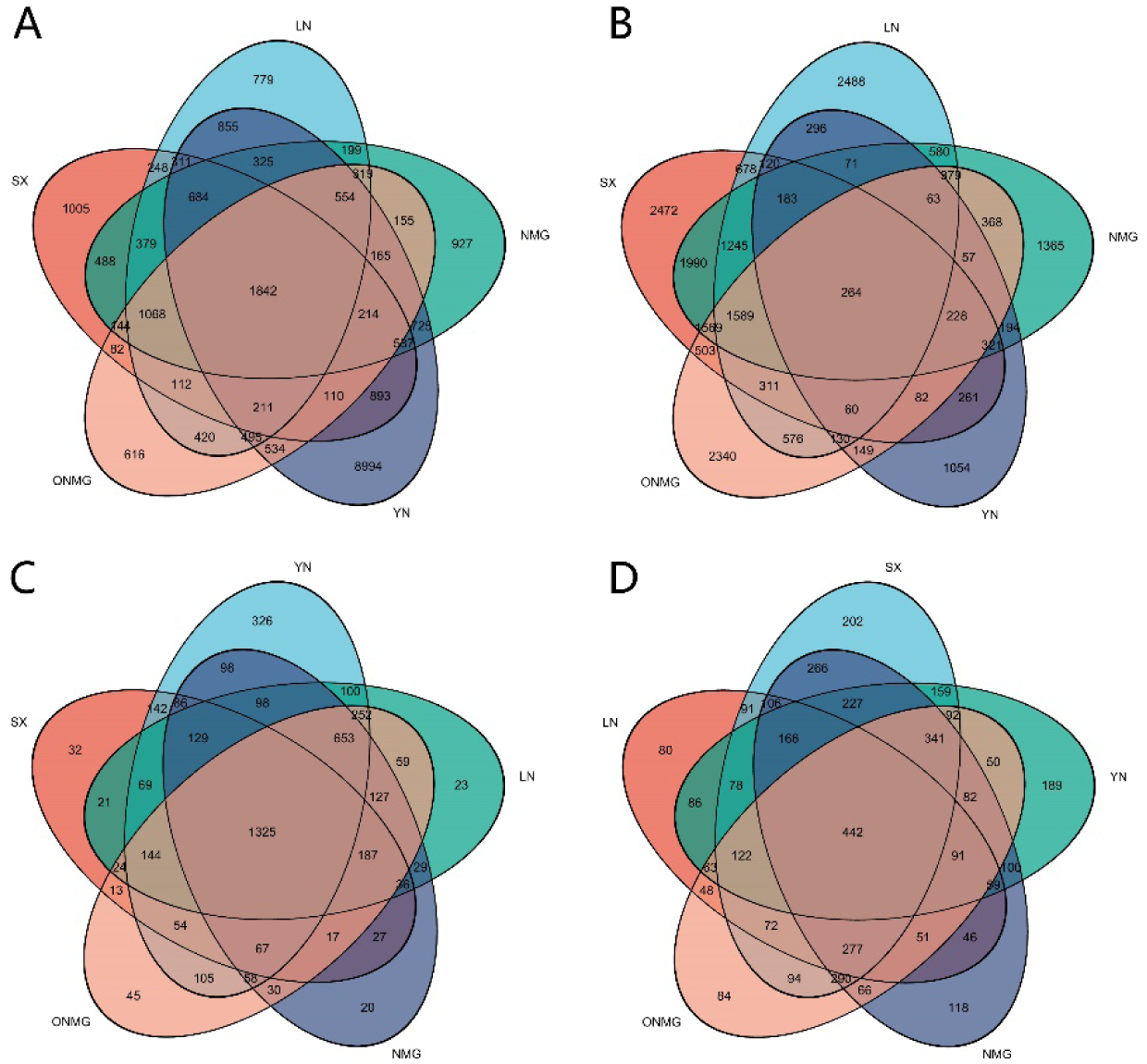
Differentially expressed genes and proteins in different diapause stages of *Lymantria dispar*. (A) Differentially expressed genes when five *L. dispar* populations enter diapause; (B) Differentially expressed genes during diapause termination in five *L. dispar* populations; (C) Differentially expressed proteins when five *L. dispar* populations enter diapause; (D) Differentially expressed proteins during diapause termination in five *L. dispar* populations.

### 3.4 Selection of Diapause-Associated Genes in Spongy Moths

The differentially expressed genes were analyzed using the KEGG pathway analysis. During the diapause initiation stage, the differentially expressed genes shared by the five populations were annotated to 214 KEGG metabolic pathways, excluding human diseases. Among these, 21 were associated with CP, 28 with EIP, 18 with GIP, 67 with metabolism, and 80 with OS. During the diapause termination stage, the differentially expressed genes shared by the five populations were annotated to 58 KEGG metabolic pathways, excluding human diseases. Among these, 11 were associated with CP, 10 with EIP, 5 with GIP, 16 with metabolism, and 16 with OS.

A total of 52 pathways were significantly enriched by differentially expressed genes during both the diapause initiation and termination stages (P < 0.05). Among these, 100 differentially expressed genes were significantly enriched in the glutathione metabolism pathway, where glutathione acts as a crucial intracellular antioxidant. In this pathway, during the diapause initiation stage, the expression of *GST* was significantly upregulated, enhancing the synthesis of glutathione S-transferase and reducing total glutathione content. However, during the diapause termination stage, the expression of *GCLC* was significantly upregulated, promoting the synthesis of glutamate–cysteine ligase and increasing the total glutathione content.

Furthermore, 70 differentially expressed genes were significantly enriched in the citrate cycle pathway, where the expression levels of *IDH1*, *IDH2*, and *icd* were markedly downregulated during the diapause initiation stage in spongy moth eggs, suppressing the activity of citrate cycle and modulating the rate of energy release.

Moreover, 40 differentially expressed genes were significantly enriched in the alanine, aspartate, and glutamate metabolism pathway, where spongy moth eggs exhibited significantly increased expressions of *GLUD1_2* and *gdhA* during the diapause initiation stage, facilitating the synthesis of mitochondrial glutamate dehydrogenase and formation of aspartate entering the urea cycle. However, during the diapause termination stage, the expression of *GOT1* was significantly upregulated, enhancing the synthesis of aspartate transaminase and leading to the formation of glutamate and oxaloacetate, thus reducing urea levels.

On the basis of the enrichment of these differentially expressed genes during the initiation and termination stages of diapause, we speculate that these genes may be involved in the diapause process of spongy moths. Thus, we next analyzed eight diapause-associated genes, namely *GST*, *GCLC*, *IDH1*, *IDH2*, *icd*, *GLUD1_2*, *gdhA*, and *GOT1*.

### 3.5 Diapause-Associated Gene Homology and Phylogenetic Analysis

The amino acid sequences of the eight selected diapause-associated genes in spongy moths were analyzed for homology by using the NCBI Protein Blast online tool. Using MEGA 7.0, the amino acid sequences of these eight diapause-associated genes were aligned with the corresponding sequences of genes from other species downloaded from NCBI. The NJ method was used for constructing a phylogenetic tree to analyze evolutionary relationships among various closely related species. Phylogenetic analysis revealed the distinct clustering of amino acid sequences encoded by each gene (Fig 6). The eight diapause-associated genes formed individual branches with genes from respective closely related species, and they were adjacent to the branches of genes from the same pathways, forming a larger cluster.

**Fig 6.**
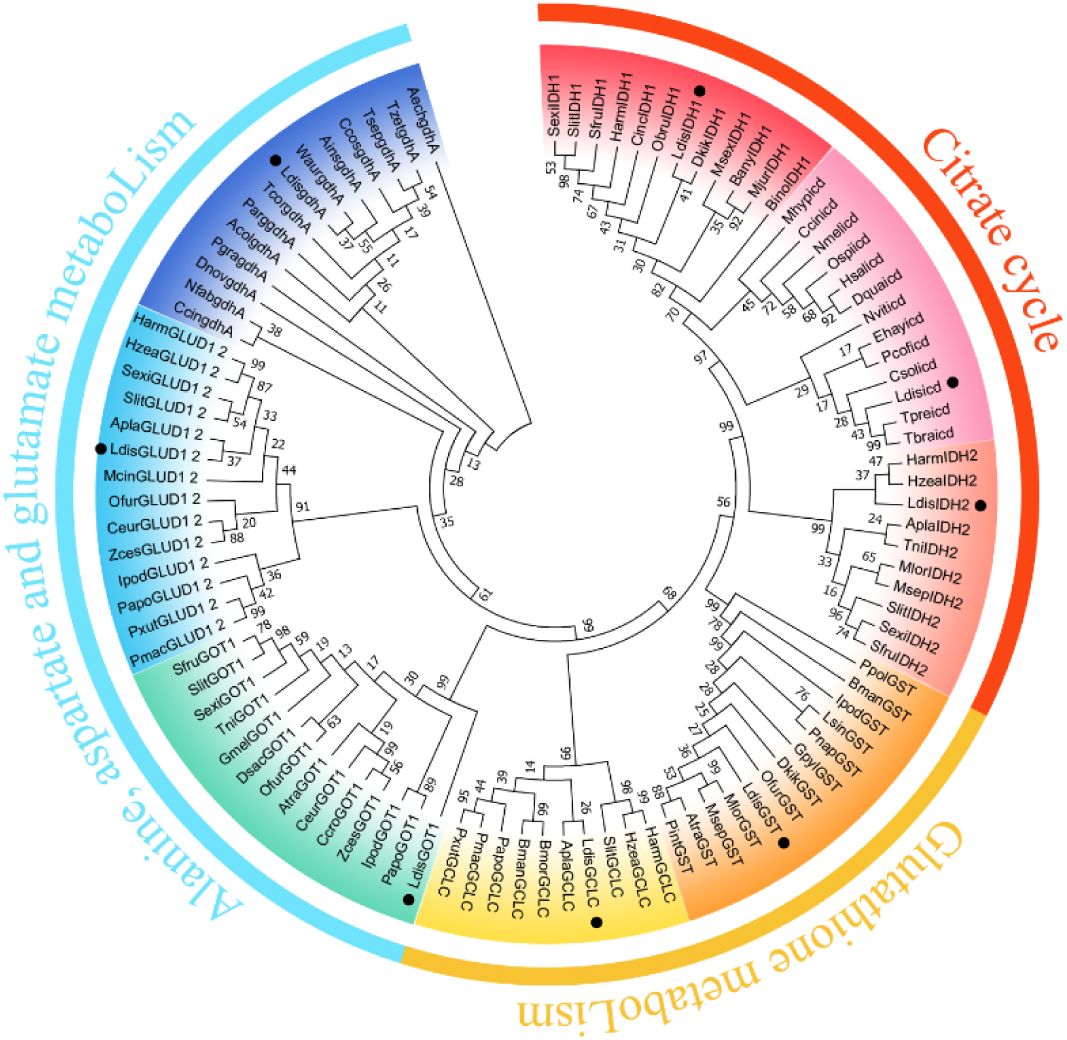
Neighbor-joining phylogenetic tree analysis of amino acid sequences of eight diapause-associated genes in *Lymantria dispar*. Black dots indicate diapause-associated genes in *L. dispar*.

### 3.6 Prediction of Three-Dimensional Structures of Diapause-Associated Proteins in Spongy Moths

The constructed structural models of diapause-associated proteins revealed that GST comprises eight α-helices and four β-strands; GCLC consists of 25 α-helices and 18 β-strands; IDH1 features 13 α-helices and 10 β-strands; IDH2 contains 12 α-helices and 10 β-strands; icd includes 11 α-helices and 12 β-strands; GLUD1_2 possesses 18 α-helices and 13 β-strands; gdhA exhibits three α-helices and a single β-strand; and GOT1 presents 13 α-helices and 10 β-strands (Fig 7).

**Fig 7.**
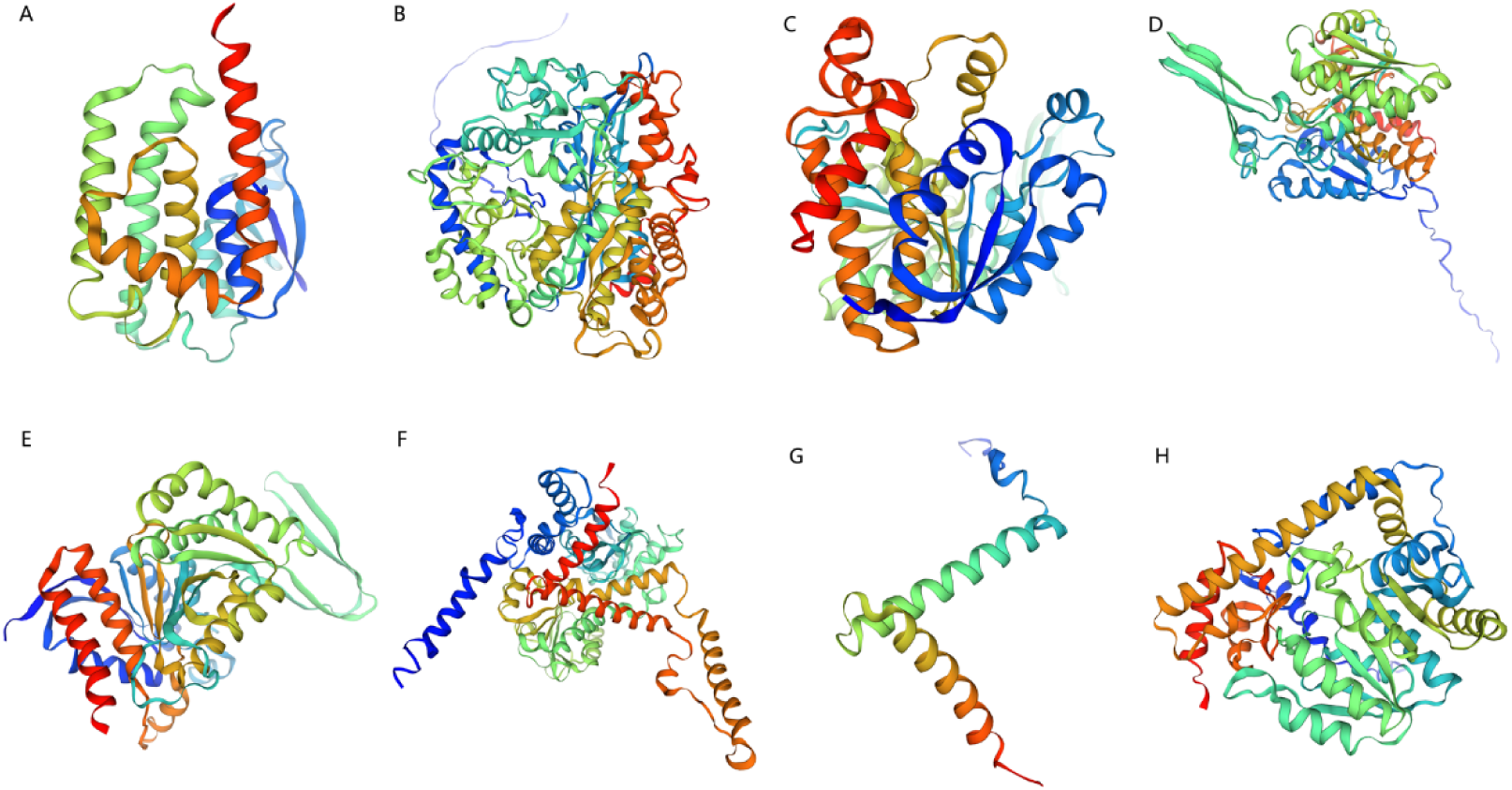
Three-dimensional protein structure prediction. GST (A), GCLC (B), IDH1 (C), IDH2 (D), icd (E), GLUD1_2 (F), gdhA (G), GOT1 (H).

The stereochemical properties of the protein models were evaluated using Ramachandran plots, which analyze the distribution of amino acids in favored, allowed, generously allowed, and disallowed conformations. For all models, over 99% of amino acid residues were positioned within the allowed regions (Fig 8). MolProbity scores were computed on the basis of modifications to the model files, including hydrogenation, followed by evaluation using clashscore, rotamers, and Ramachandran criteria. Lower MolProbity scores indicated higher model quality, with scores annotated in green (good), yellow (caution), and red (warning) (Table 3). For more information on table colors and cutoffs, see: http://molprobity.biochem.duke.edu/help/validation_options/summary_table_guide.html.

**Fig 8.**
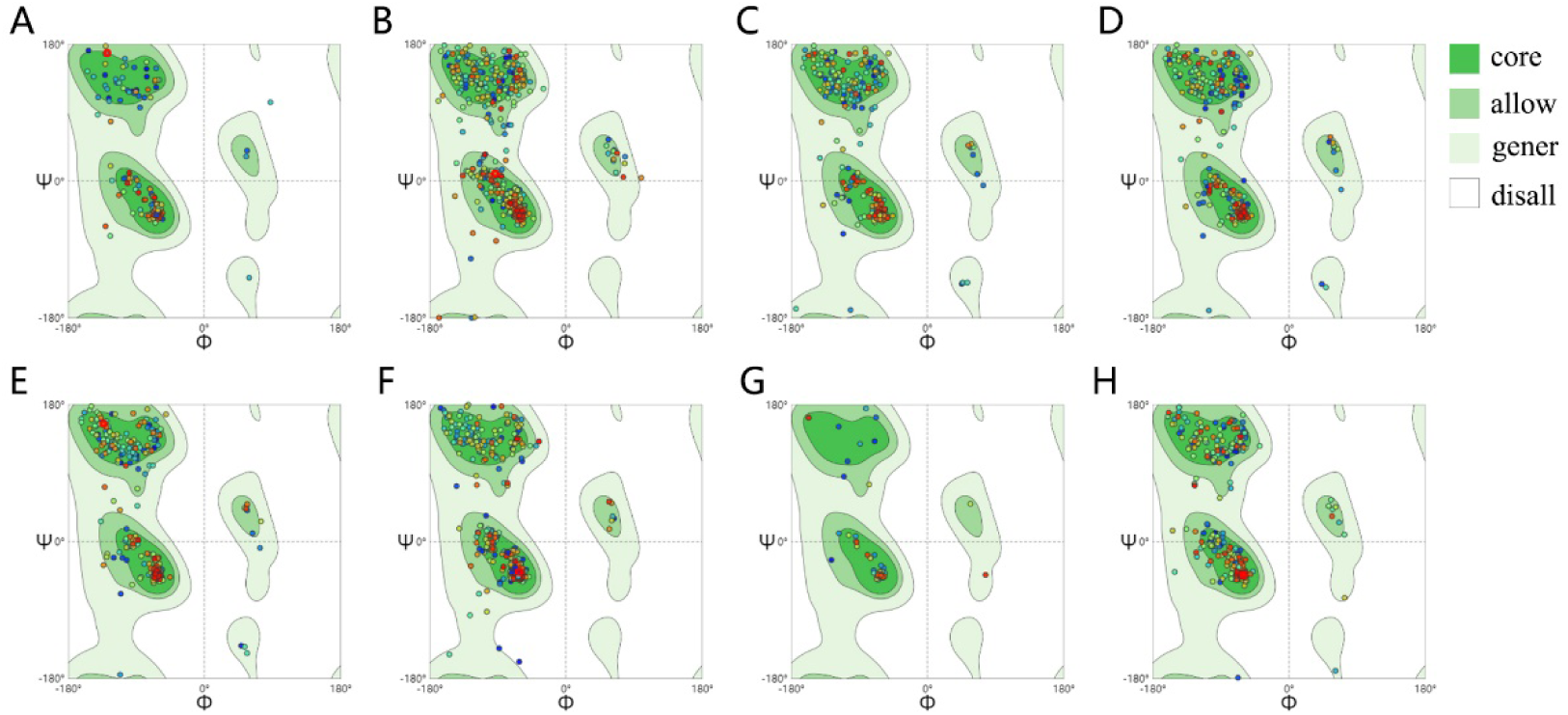
Ramachandran plots verification of three-dimensional protein structure prediction. GST (A), GCLC (B), IDH1 (C), IDH2 (D), icd (E), GLUD1_2 (F), gdhA (G), GOT1 (H).

**Table 3.** Evaluation of the three-dimensional protein structure prediction model.

| Model | Clashscore | Favored rotamers | Ramachandran favored | MolProbity score |
| --- | --- | --- | --- | --- |
| GST | 1.43 | 96.28% | 98.16% | 0.88 |
| GCLC | 0.95 | 96.30% | 93.83% | 1.20 |
| IDH1 | 0.61 | 98.01% | 95.81% | 0.99 |
| IDH2 | 0.15 | 98.38% | 95.35% | 0.88 |
| icd | 0.34 | 98.39% | 95.89% | 0.91 |
| GLUD1_2 | 0.71 | 96.89% | 96.51% | 0.96 |
| gdhA | 0 | 100.00% | 96.05% | 0.77 |
| GOT1 | 1.24 | 97.98% | 98.28% | 0.84 |

Clashscore is the number of serious steric overlaps (>0.4 Å) per 1000 atoms. MolProbity score combines the clashscore, rotamer, and Ramachandran evaluations into a single score, normalized to be on the same scale as X-ray resolution. Green: good; Yellow: caution; Red: warning.

## 4. Discussion

### 4.1 Glutathione Metabolism Pathway

Glutathione exists in the reduced glutathione and oxidized glutathione (GSSG) forms. Among these, reduced glutathione, the most crucial intracellular antioxidant, plays a critical role in clearing intracellular peroxides and free radicals and repairing damaged tissues, thereby protecting mitochondria from oxidative stress generated during physiological and pathological processes [33]. The activity of catalase, glutathione S-transferase, and dehydroascorbate reductase significantly differs between the mitochondria of diapausing and non-diapausing larvae of the European corn borer [34]. In the diapause process of *Chlosyne lacinia*, glutathione transferase and isocitrate dehydrogenase-NADP(+) play a role in responding to oxidative stress [35]. Hydrogen peroxide metabolism is closely associated with diapause regulation in silkworms [36], because hydrogen peroxide can promote cell division [37]. Exogenous application of hydrogen peroxide can inhibit or alleviate diapause [38]. The glutathione redox cycle plays a crucial role in the clearance of hydrogen peroxide [39]. Changes in the expression levels of glutathione during different diapause stages may be related to the process of hydrogen peroxide clearance.

During the onset of diapause in spongy moth eggs, the expression of GST in the glutathione metabolism pathway is upregulated, leading to a decrease in total glutathione content. By contrast, upon diapause termination, the expression of GCLC is upregulated, leading to an increase in total glutathione content. Thus, hydrogen peroxide may also have a regulatory role in inhibiting diapause in spongy moths.

### 4.2 Citric Acid Cycle Pathway

Isocitrate dehydrogenase (IDH) is a key enzyme that regulates the rate of the tricarboxylic acid (TCA) cycle. IDH catalyzes the oxidative decarboxylation of isocitrate to α-ketoglutarate, serving as an important source of cellular NADPH [40]. NADPH plays a crucial role in counteracting intracellular oxidative stress by reducing oxidized glutathione [41] and serves as an electron donor in numerous enzyme reactions, biosynthesis pathways, and detoxification processes [42]. An increase in NADPH content indicates that the synthesis of NADP exceeds its degradation, suggesting that increased cellular respiration during cell division requires substantial NADPH for biosynthesis and antioxidation. A study reported that the IDH content in nondiapause B. mori eggs is significantly higher than that in diapause eggs [43].

Upon the initiation of diapause, spongy moth eggs exhibit a decrease in the expression of IDH1, IDH2, and icd in the citric acid cycle, inhibiting this cycle. The citric acid cycle is a common metabolic pathway for energy acquisition in aerobic organisms. During insect diapause, the citric acid cycle is inhibited [44–46], reducing the overall reaction rate. Our study findings indicate that spongy moths exhibit a state of low-energy metabolism during diapause, aligning with the result of a previous study that demonstrated a significant decrease in metabolic levels when insects enter diapause stage [47].

Consistent with our results, studies on insects, such as *Sitodiplosis mosellana* [48] and *Coccinella septempunctata* [49], have reported a significant downregulation of IDH during diapause, indicating the inhibition of the TCA cycle. However, other studies have found that in some species, such as *C. lacinia* [35] and *Exorista civilis* [50], IDH content significantly increases during diapause, suggesting a potential role of IDH in response to oxidative stress [51].

### 4.3 Alanine, Aspartate, and Glutamate Metabolism Pathway

Glutamate plays diverse physiological roles and is a key neurotransmitter. In vertebrates, glutamate primarily functions by promoting calcium ion transport to facilitate excitatory neurotransmission in neuronal differentiation, migration, and survival [52]. In invertebrates, glutamate can serve as both an excitatory and inhibitory neurotransmitter. Moreover, glutamate is a component of proteins and peptides and a vital binding component for glutathione synthesis [53]. Phosphoenolpyruvate carboxykinase (PEPCK) plays a central role in mediating insect responses to environmental stress and diapause processes [54, 55]. PEPCK is the main regulatory enzyme of glyceroneogenesis, and this process requires precursors, such as alanine and glutamine, to synthesize glycerol-3-phosphate or triacylglycerol. Moreover, aspartate and glutamate are crucial amino acids in the urea synthesis process. Mitochondrial glutamate dehydrogenase removes the amino group from glutamate, converting it under enzymatic action into carbamoyl phosphate, which enters the urea cycle. Aspartate in the urea cycle is produced from glutamate and oxaloacetate, catalyzed by aspartate aminotransferase. Zhang et al. demonstrated that *H. armigera* accumulates large amounts of urea to withstand low temperatures during diapause [56].

In spongy moth eggs, upon diapause initiation, the expression of GLUD1_2 and gdhA increases in the alanine, aspartate, and glutamate metabolism pathway. This alteration leads to decreased glutamate levels, increased aspartate levels, and the activation of the urea cycle. Upon diapause termination, GOT1 expression is upregulated, resulting in increased glutamate levels and decreased aspartate levels, thereby inhibiting the urea cycle. These changes suggest that spongy moths experience a reduction in the metabolic rate upon entering diapause, which inhibits amino acid synthesis and leads to excess ammonia, which is diverted into the urea cycle. By contrast, diapause termination is associated with the increased metabolic activity and urea cycle inhibition. In addition, spongy moths may exhibit enhanced cold tolerance by accumulating excessive urea during diapause.

### 4.4 Limitations and Future Directions

The finding of this study suggest that diapause-associated genes improve cold tolerance and metabolism in spongy moths during diapause. Changes in the expression levels of diapause-associated genes regulate the onset of diapause. Previous studies have revealed that FSMC requires less time to break diapause under low temperatures than ESM, and the intensity of diapause is also lower [11]. However, this study focused on only four populations of FSMC and one hybrid population of FSMC and ESM due to the lack of ESM sources. Thus, research on diapause genes in ESM is required. When ESM enter diapause, whether the molecular mechanism of diapause in ESM is the same as that in FSMC, and whether differences exist in the expression patterns of the eight diapause-associated genes remain to be determined. Subsequent studies should focus on the diapause molecular mechanisms in ESM and conduct a comparative analysis of diapause-associated genes between ESM and FSMC as well as explore whether the reasons for the faster breaking of diapause in FSMC can be explained from a genetic perspective.

## Acknowledgments

We would like to thank Xiaoxiao Chang (Beijing Forestry University) for technical guidance, Yixin Qiu (Beijing Forestry University), Yanyi Lu (Beijing Forestry University) and Yuxuan Li (Beijing Forestry University) for assistance in raising spongy moth.

## Supporting information

**S1 File. NR Annotation Information Table-Transcriptome functional annotation.**

**S2 File. NR Annotated Species Distribution Table-Transcriptome functional annotation.**

**S3 File. Swiss-Prot Annotation Information Table-Transcriptome functional annotation.**

**S4 File. Pfam Annotation Information Table-Transcriptome functional annotation;.**

**S5 File. EggNOG Classification Statistics Table-Transcriptome functional annotation.**

**S6 File. GO Classification Statistics Table-Transcriptome functional annotation.**

**S7 File. KEGG Classification Statistics Table-Transcriptome functional annotation.**

**S8 File. Pfam Annotation Information Table-Proteome functional annotation.**

**S9 File. EggNOG Classification Statistics Table-Proteome functional annotation.**

**S10 File. GO Classification Statistics Table-Proteome functional annotation.**

**S11 File. KEGG Classification Statistics Table-Proteome functional annotation.**

**S12 File. Subloc Annotation Information Table-Proteome functional annotation.**

**S13 File. GO Enrichment Analysis Statistical Table of DEGs upon Diapause Initiation.**

**S14 File. GO Enrichment Analysis Statistical Table of DEGs upon Diapause Termination.**

**S15 File. KEGG Enrichment Analysis Statistical Table of DEGs upon Diapause Initiation.**

**S16 File. KEGG Enrichment Analysis Statistical Table of DEGs upon Diapause Termination.**

## References

1. Boukouvala MC, Kavallieratos NG, Skourti A, Pons X, Alonso CL, Eizaguirre M, Fernandez EB, Solera ED, Fita S, Bohinc T, Trdan S, Agrafioti P and Athanassiou CG, Lymantria dispar (L) (Lepidoptera: Erebidae): Current Status of Biology, Ecology, and Management in Europe with Notes from North America. Insects 13:854 (2022). 10.3390/insects13090854.

2. Keena MA, Côté MJ, Grinberg PS and Wallner WE, World Distribution of Female Flight and Genetic Variation in Lymantria dispar (Lepidoptera: Lymantriidae). Environ. Entomol. 37:636–649 (2008). 10.1603/0046-225x(2008)37[636:Wdoffa]2.0.Co;2.

3. Keena MA and Richards JY, Comparison of Survival and Development of Gypsy Moth Lymantria dispar L. (Lepidoptera: Erebidae) Populations from Different Geographic Areas on North American Conifers. Insects 11:260 (2020). 10.3390/insects11040260.

4. Kim MJ, Kim KE, Lee CY, Park Y, Jung JK and Nam Y, Effect of Chilling Temperature on Survival and Post-Diapause Development of Korean Population of Lymantria dispar asiatica (Lepidoptera: Erebidae) Eggs. Forests 13:2117 (2022). 10.3390/f13122117

5. Djoumad A, Nisole A, Zahiri R, Freschi L, Picq S, Gundersen-Rindal DE, Sparks ME, Dewar K, Stewart D, Maaroufi H, Levesque RC, Hamelin RC and Cusson M, Comparative analysis of mitochondrial genomes of geographic variants of the gypsy moth, Lymantria dispar, reveals a previously undescribed genotypic entity. Sci. Rep. 7:14245 (2017). 10.1038/s41598-017-14530-6.

6. Keena MA, Grinberg PS and Wallner WE, Inheritance of female flight in Lymantria dispar (Lepidoptera: Lymantriidae). Environ. Entomol. 36:484–494 (2007). 10.1603/0046-225X(2007)36[484:IOFFIL]2.0.CO;2.

7. Sandquist RE, Richerson JV and Cameron EA, Flight of North American Female Gypsy Moths. Environ. Entomol. 2:957–958 (1973). 10.1093/ee/2.5.957.

8. Koshio C, Pre-ovipositional Behaviour of the Female Gypsy Moth, Lymantria dispar L. (Lepidoptera, Lymantriidae). Appl. Entomol. Zool. 31:1–10 (1996). 10.1016/j.jnoncrysol.2004.08.132.

9. Schaefer WP, Weseloh MR, Sun XL, Wallner EW and Yan JJ, Gypsy Moth, Lymantria (=Ocneria) dispar (L.) (Lepidoptera: Lymantriidae), in the People’s Republic of China. Environ. Entomol. 13:1535–1541 (1984).

10. Wallner WE, Humble LM, Levin RE, Baranchikov YN and Carde RT, Response of Adult Lymantriid Moths to Illumination Devices in the Russian Far East. J. Econ. Entomol. 88:337–342 (1995). 10.1093/jee/88.2.337.

11. Keena MA, Comparison of the Hatch of Lymantria dispar (Lepidoptera: Lymantriidae) Eggs from Russia and the United States after Exposure to Different Temperatures and Durations of Low Temperature. Ann. Entomol. Soc. Am. 89:564–572 (1996). 10.1093/aesa/89.4.564.

12. Taylor F, Insect life histories: seasonal adaptations of insects. Science 232:1152 (1986). 10.1126/science.232.4754.1152-a.

13. Gray DR, Logan JA, William RF and Carlson JA, Toward a Model of Gypsy Moth Egg Phenology: Using Respiration Rates of Individual Eggs to Determine Temperature-Time Requirements of Prediapause Development. Environ. Entomol. 20:1645–1652 (1991). 10.1093/ee/20.6.1645.

14. Li Y, The study on the factors to break Diapause and the Change of several Macronutrient Matrials in Asian Gypsy Moth, Lymantria dispar asiatica. Beijing Forestry University (2014).

15. Gray DR, Ravlin FW, Régnière J and Logan JA, Further advances toward a model of gypsy moth (Lymantria dispar (L.)) egg phenology: Respiration rates and thermal responsiveness during diapause, and age-dependent developmental rates in postdiapause. J. Insect Physiol. 41:247–256 (1995). 10.1016/0022-1910(94)00102-M.

16. Kostál V, Eco-physiological phases of insect diapause. J. Insect Physiol. 52:113–127 (2006). 10.1016/j.jnoncrysol.2004.08.132.

17. Wei J, Luo YQ, Shi J, Wang DP and Shen SW, Impact of temperature on postdiapause and diapause of the Asian gypsy moth, Lymantria dispar asiatica. J. Insect Sci. 14:5 (2014). 10.1673/031.014.5.

18. Keena MA, Inheritance and World Variation in Thermal Requirements for Egg Hatch in Lymantria dispar (Lepidoptera: Erebidae). Environ. Entomol. 45:1–10 (2015). 10.1093/ee/nvv163.

19. Ponomarev VI, Klobukov GI, Ilyinykh AV and Dubovskiy IM, Adaptation Features of Diapause Duration of the Gypsy Moth Lymantria dispar (L.) from Populations of Different Latitudinal Origination. Contemp. Probl. Ecol. 12:1–9 (2019). 10.1134/S1995425519010098.

20. Lee KY and Denlinger DL, A Role for Ecdysteroids in the Induction and Maintenance of the Pharate First Instar Diapause of the Gypsy Moth, Lymantria dispar. J. Insect Physiol. 43:289–296 (1997). 10.1016/S0022-1910(96)00082-0.

21. Atay-Kadiri Z and Benhsain N, The diapause of gypsy moth, Lymantria dispar (L.) (Lepidoptera: Lymantriidae). Trends Comp. Endocrinol. Neurobiol. 1040:219–223 (2005). 10.1196/annals.1327.028.

22. Li Y, Liang Y and Shi J, The Change of Several lmportant Macronutrient Matrials in Asian Gypsy Moth, Lymantria dispar (Lepidoptera: Lymantriidae). Chin. Agric. Sci. Bull. 30:55–60 (2014).

23. Gu SH, Lin PL and Chang CH, Expressions of sugar transporter genes during Bombyx mori embryonic development. J. Exp. Zool. Part A 339:788–798 (2023). 10.1002/jez.2729.

24. Gong J, Zhang W, Wang QL, Zhu ZJ, Pang JX and Hou Y, Genome-wide identification of the BmAKR gene family in the silkworm (Bombyx mori) and their expression analysis in diapause eggs and nondiapause eggs. Chin. J. Biotechnol. 39:4982–4995 (2023). 10.13345/j.cjb.230105.

25. Chen YR, Jiang T, Zhu J, Xie YC, Tan ZC, Chen YH, Tang SM, Hao BF, Wang SP, Huang JS and Shen XJ, Transcriptome sequencing reveals potential mechanisms of diapause preparation in bivoltine silkworm Bombyx mori (Lepidoptera: Bombycidae). Comp. Biochem. Physiol. D-Genomics Proteomics 24:68–78 (2017). 10.1016/j.cbd.2017.07.003.

26. Reynolds JA, Nachman RJ and Denlinger DL, Distinct microRNA and mRNA responses elicited by ecdysone, diapause hormone and a diapause hormone analog at diapause termination in pupae of the corn earworm, Helicoverpa zea. Gen. Comp. Endocrinol. 278:68–78 (2019). 10.1016/j.ygcen.2018.09.013.

27. Chen LZ, Ma WH, Wang XP, Niu CY and Lei CL, Analysis of pupal head proteome and its alteration in diapausing pupae of Helicoverpa armigera. J. Insect Physiol. 56:247–252 (2010). 10.1016/j.jinsphys.2009.10.008.

28. Lu Q, Li Y, Liao J, Ni ZH, Xia SC, Yang MF, Li HY and Guo JJ, Histone acetylation is associated with pupal diapause in cotton bollworm, Helicoverpa armigera. Pest Manage. Sci. 80:1400–1411 (2024). 10.1002/ps.7870.

29. Lebenzon JE, Torson AS and Sinclair BJ, Diapause differentially modulates the transcriptomes of fat body and flight muscle in the Colorado potato beetle. Comp. Biochem. Physiol. D-Genomics Proteomics 40:100906 (2021). 10.1016/j.cbd.2021.100906.

30. Dong CL, Lu MX and Du YZ, Transcriptomic analysis of pre-diapause larvae of Chilo suppressalis (Walker) (Lepidoptera: Pyralidae) in natural populations. Comp. Biochem. Physiol. D-Genomics Proteomics 40:100903 (2021). 10.1016/j.cbd.2021.100903.

31. Wang YJ, Structural Anatomy and Internal Regulation Mechanism Related to the Regulation of Diapause by the Gypsy Moth. Beijing Forestry University (2022).

32. Bradford MM, A rapid and sensitive method for the quantitation of microgram quantities of protein utilizing the principle of protein-dye binding. Anal. Biochem. 72:248–254 (1976). 10.1016/0003-2697(76)90527-3.

33. Hutter DE, Till BG and Greene JJ, Redox state changes in density-dependent regulation of proliferation. Exp. Cell Res. 232:435–438 (1997). 10.1006/excr.1997.3527.

34. Jovanovic-Galovic A, Blagojevic DP, Grubor-Lajsic G, Worland MR and Spasic MB, Antioxidant Defense in mitochondria during diapause and postdiapause development of European corn borer (Ostrinia nubilalis, Hubn.). Arch. Insect Biochem. Physiol. 64:111–119 (2007). 10.1002/arch.20160.

35. Moreira DC, Paula DP and Hermes-Lima M, Changes in metabolism and antioxidant systems during tropical diapause in the sunflower caterpillar Chlosyne lacinia (Lepidoptera: Nymphalidae). Insect Biochem. Mol. Biol. 134:103581 (2021). 10.1016/j.ibmb.2021.103581.

36. Zhao LC, Sima YH, Shen XM, Metabolism of Hydroen peroxide in the Course of Embryonic Development in Silkworm. Dev. Reprod. Biol. 8:41–48 (1999).

37. Burdon RH, Gill V and Rice-Evans C, Oxidative Stress and Tumour Cell Proliferation. Free Radical Res. Commun. 11:65–76 (1990). 10.3109/10715769009109669.

38. Robbins HM, Van Stappen G, Sorgeloos P, Sung YY, MacRae TH and Bossier P, Diapause termination and development of encysted Artemia embryos: roles for nitric oxide and hydrogen peroxide. J. Exp. Biol. 213:1464–1470 (2010). 10.1242/jeb.041772.

39. Forman HJ, Zhang HQ, Rinna A, Glutathione: Overview of its protective roles, measurement, and biosynthesis. Mol. Asp. Med. 30:1–12 (2009). 10.1016/j.mam.2008.08.006.

40. Fantin VR, Dang L, White DW, Gross S, Bittinger MA, Driggers EM, Jang HG, Jin SF, Keenan MC, Marks KM, Yen KE, Ward PS, Prins RM, Liau LM, Bennett BD, Rabinowitz JD, Cantley LC, Thompson CB, Heiden MV and Su SM, Cancer-associated IDH1 mutations produce 2-hydroxyglutarate. Cancer Res. 70:33 (2010). 10.1158/1538-7445.AM10-33.

41. Nicholas JK and Antje VS, The oxidative pentose phosphate pathway: structure and organisation. Curr. Opin. Plant Biol. 6:236–246 (2003). 10.1016/S1369-5266(03)00039-6.

42. Leterrier M, Barroso JB, Palma JM and Corpas FJ, Cytosolic NADP-isocitrate dehydrogenase in Arabidopsis leaves and roots. Biol. Plant. 56:705–710 (2012). 10.1007/s10535-012-0244-6.

43. Yao JM, Wan HX, Sima YH and Zhao LC, Instant hydrochloride acid soaking enhances NAD(H) and NADP(H) levels in diapausing eggs of the silkworm, Bombyx mori. Acta Entomol. Sin. 55:24–28 (2012). 10.16380/j.kcxb.2012.01.003.

44. Michaud MR and Denlinger DL, Shifts in the carbohydrate, polyol, and amino acid pools during rapid cold-hardening and diapause-associated cold-hardening in flesh flies (Sarcophaga crassipalpis): a metabolomic comparison. J. Comp. Physiol. B-Biochem. Syst. Environ. Physiol. 177:753–763 (2007). 10.1007/s00360-007-0172-5.

45. Xu WH, Lu YX and Denlinger DL, Cross-talk between the fat body and brain regulates insect developmental arrest. Proc. Natl. Acad. Sci. U. S. A. 109:14687–14692 (2012). 10.1073/pnas.1212879109.

46. Lu YX, Zhang Q and Xu WH, Global Metabolomic Analyses of the Hemolymph and Brain during the Initiation, Maintenance, and Termination of Pupal Diapause in the Cotton Bollworm, Helicoverpa armigera. PLoS One 9:e99948 (2014). 10.1371/journal.pone.0099948.

47. Ragland GJ, Fuller J, Feder JL and Hahn DA, Biphasic metabolic rate trajectory of pupal diapause termination and post-diapause development in a tephritid fly. J. Insect Physiol. 55:344–350 (2009). 10.1016/j.jinsphys.2008.12.013.

48. Gong ZJ, Wu YQ, Miao J, Duan Y, Jiang YL and Li T, Global Transcriptome Analysis of Orange Wheat Blossom Midge, Sitodiplosis mosellana (Gehin) (Diptera: Cecidomyiidae) to Identify Candidate Transcripts Regulating Diapause. PLOS ONE 8:e71564 (2013). 10.1371/journal.pone.0071564.

49. Ren XY, Zhang LS, Han YH, An T, Liu Y, Li YY and Chen HY, Proteomic research on diapause-related proteins in the female ladybird, Coccinella septempunctata L. Bull. Entomol. Res. 106:168–174 (2016). 10.1017/S0007485315000954.

50. Zhang B, Han HB, Xu LB, Li YR, Song MX and Liu AP, Transcriptomic analysis of diapause-associated genes in Exorista civilis Rondani (Diptera: Tachinidae). Arch. Insect Biochem. Physiol. 107:e21789 (2021). 10.1002/arch.21789.

51. Niazi AK, Bariat L, Riondet C, Carapito C, Mhamdi A, Noctor G and Reichheld JP, Cytosolic Isocitrate Dehydrogenase from Arabidopsis thaliana Is Regulated by Glutathionylation. Antioxidants 8:16 (2019). 10.3390/antiox8010016.

52. Torii K, Uneyama H and Nakamura E, Physiological roles of dietary glutamate signaling via gut-brain axis due to efficient digestion and absorption. J. Gastroenterol. 48:442–451 (2013). 10.1007/s00535-013-0778-1.

53. Colowick NP, Kaplan NP and Meister A, Glutamate, glutamine, glutathione, and related compounds. Methods Enzymol. 113:1–723 (1985).

54. Poelchau MF, Reynolds JA, Denlinger DL, Elsik CG and Armbruster PA, A de novo transcriptome of the Asian tiger mosquito, Aedes albopictus, to identify candidate transcripts for diapause preparation. BMC Genomics 12:619 (2011). 10.1186/1471-2164-12-619.

55. Ragland GJ, Egan SP, Feder JL, Berlocher SH and Hahn DA, Developmental trajectories of gene expression reveal candidates for diapause termination: a key life-history transition in the apple maggot fly Rhagoletis pomonella. J. Exp. Biol. 214:3948–3959 (2011). 10.1242/jeb.061085.

56. Zhang Q, Lu YX and Xu WH, Proteomic and metabolomic profiles of larval hemolymph associated with diapause in the cotton bollworm, Helicoverpa armigera. BMC Genomics 14:751 (2013). 10.1186/1471-2164-14-751.

